# Determination of insulin secretion from stem cell-derived islet organoids with liquid chromatography-tandem mass spectrometry

**DOI:** 10.1101/2022.10.24.512117

**Authors:** Christine Olsen, Chencheng Wang, Shadab Abadpour, Elsa Lundanes, Audun Skau Hansen, Frøydis Sved Skottvoll, Hanne Scholz, Steven Ray Wilson

## Abstract

Organoids are laboratory-grown 3D organ models, mimicking human organs for e.g. drug development and personalized therapy. Islet organoids (typically 100-200 μm), which can be grown from the patient’s own cells, are emerging as prototypes for transplantation-based therapy of diabetes. Selective methods for quantifying insulin production from islet organoids are needed, but sensitivity and carry-over have been major bottlenecks in previous efforts. We have developed a reverse phase liquid chromatography-tandem mass spectrometry (RPLC-MS/MS) method for studying the insulin secretion of islet organoids. In contrast to our previous attempts using nano-scale LC columns, conventional 2.1 mm inner diameter LC column (combined with triple quadrupole mass spectrometry) was well suited for sensitive and selective measurements of insulin secreted from islet organoids with low microliter-scale samples. Insulin is highly prone to carry-over, so standard tubings and injector parts were replaced with shielded fused silica nanoViper™ connectors. As samples were expected to be very limited, an extended Box-Behnken experimental design for the MS settings was conducted to maximize performance. The finale method has excellent sensitivity, accuracy and precision (limit of detection: ≤ 0.2 pg/μL, relative error: ≤ ±10%, relative standard deviation: < 10%), and was well suited for measuring 20 μL amounts of Krebs buffer containing insulin secreted from islet organoids.

## Introduction

Hormonal imbalance, i.e. endocrine disorders (ED), is typically treated with lifelong daily hormone replacement therapy (HRT). However, HRT is limited in achieving physiological mimicry, e.g. in type 1 diabetes (T1D) patients, where injections of exogenous insulin do not match the precise control over the blood glucose as that achieved by the insulin-producing beta cells (found in pancreatic islets) in healthy individuals, leading to incidents of hyper- and hypoglycemia [1, 2]. An alternative is beta cell replacement either of donor-derived solid pancreas or isolated islets transplantation, but the approach is adversely affected by the scarcity of donors and the use of immunosuppressive medications due to risk of rejection [1, 3]. Ex vivo generated islet organoids is largely thought to be one possible solution to an unlimited source and achieve treatment of T1D patients [3-7]. However, protocols for generating organoids (i.e. laboratory-grown 3D organ models) are still under development, and there is a need for analytical tools for studying the traits and dynamics of organoids, including stem cell-derived islet (SC-islet) organoids [8].

Renewable in-vitro-produced SC-islets provide virtually unlimited cell resources for diabetes transplantation-based therapy. A critical feature to characterize SC-islet is their ability to secrete insulin in response to glucose [7, 9]. Insulin secretion is typically measured with enzyme-linked immunosorbent assay (ELISA), but ELISA methods have selectivity weaknesses, i.e. inability to distinguish insulin from structural analogs [10].

Our research group focuses on applying liquid chromatography-mass spectrometry (LC-MS) to study the traits of various organoids [11-14], and we here describe the first MS-based method for highly selective monitoring of insulin excreted from islet organoids. Main ingredients of the method were: The use of conventional-sized columns rather than columns in the nano-format, which we previously found were prone to carry-over and surprisingly unsatisfactory sensitivity for islet organoids [15]; Exploring tubing and injection hardware for reduced carry-over of insulin, which is highly prone to non-defined adsorption [15, 16]; Applying experimental design to maximize mass spectrometric sensitivity for very limited samples. As a proof-of-concept, we present findings concerning the determination of insulin secretion in SC-islets incubated in Krebs buffer with various levels of glucose and KCl, which stimulate insulin secretion [17].

## 1 Materials and Methods

### 1.1 Chemicals

Acetonitrile (ACN, LC-MS grade), bovine serum albumin (BSA, ≥ 98%), dimethyl sulfoxide (DMSO, ≥ 99.7%), formic acid (FA, 98%), insulin from bovine pancreas (HPLC grade), and recombinant insulin human (≥ 98%) were all purchased from Sigma-Aldrich. Water (LC-MS grade), and methanol (MeOH, LC-MS grade) were obtained from VWR Chemicals (Radnor, PA, USA). Gibco™ basal cell medium MCDB131, GlutaMAX™ supplement (cat. no. 35050061) and minimum essential medium non-essential amino acids (MEM NEAA) stock solution by Gibco™ was acquired from Thermo Fisher Scientific (Waltham, MA, USA). Krebs buffer was prepared in-house and consists of the following chemicals of analytical grade: 10mM HEPES, 128mM NaCl, 5mM KCl, 2.7mM CaCl_2_, 1.2mM MgSO_4_, 1mM Na_2_HPO_4_, 1.2mM KH_2_PO_4_, 5mM NaHCO_3_, and 0.1% BSA.

### 1.2 Preparation of islet maturation cell medium

Islet maturation cell medium was prepared in-house by adding 1% Penicillin/Streptavidin (Pen/Strep), 2.5 mM of glucose (final concentration in cell medium being 8 mM of glucose), 2% BSA, 10 μg/mL of heparin, 1 μM of ZnSO_4_, 1% of GlutaMAX™ stock solution and 1% of MEM NEAA stock solution to basal cell medium MCDB131 stock solution.

### 1.3 Preparation of human and bovine insulin solutions

Aqueous water solutions of human and bovine insulin (internal standard) were prepared individually by dissolving 1 mg of insulin powder in 1 mL of 0.1% FA in water. All solutions containing proteins were prepared in protein low binding tubes from Sarstedt (Nümbrecht, Germany). The 1 mg/mL stock solutions were further diluted to working solutions consisting of 10 ng/μL human or bovine insulin, divided into 100 μL aliquots, and kept at -20 °C until use or for a maximum of three months. Separate standard solutions of 10 ng/μL of human and bovine insulin in a 1+1 mixture of ACN and water were prepared for direct injections on the MS. For assessment of the LC method (method evaluation solutions), the working solution of human insulin was further diluted to 125 pg/μL with water and spiked with 2 ng/μL of bovine insulin solution to give a concentration of 125 pg/μL of bovine insulin.

Working and method evaluation solutions with human and bovine insulin in islet maturation cell medium and Krebs buffer were prepared in the same manner as described for water-based solutions, with the exception being the amount of FA; 1% FA in cell medium and 0.5% FA in Krebs buffer.

#### 1.3.1 Preparation of calibration standard solutions and quality controls

Calibration standard solutions and quality controls (QC) were prepared by mixing freshly thawed working solutions of human and bovine insulin and diluted to appropriate concentrations. Calibration standard solutions in 1.0% FA cell medium consisted of [7.8, 15.6, 31.25, 62.5, 125.0, and 250.0] pg/μL human insulin with 125.0 pg/μL bovine insulin. Calibration standard solutions in 0.5% FA Krebs buffer consisted of [0.2, 0.5, 1.0, 3.0, 5.0, and 10.0] pg/μL human insulin with 5.0 pg/μL bovine insulin.

### 1.4 Cell culture and differentiation

SC-islets were generated from human pluripotent cell line H1 (WA01, WiCell, Madison, WI, USA). Undifferentiated H1 cells were cultured in Essential 8™ Medium (Gibco) on tissue culture plates coated with Geltrex™ in a humidified incubator containing 5% CO_2_ at 37 °C. Undifferentiated cells were passaged with 0.5 mM EDTA. To initiate differentiation, undifferentiated cells were seeded at 2×10^5^ cells/cm^2^ in Geltrex-coated cell culture plates. The subsequent stepwise differentiation schedules and media recipes are provided in **SI-Table 1**. On day 7 of stage 6, the differentiated cells were dissociated with TrypLE and seeded at 1 × 10^6^ cells/mL in ultra-low attachment cell culture plates from Corning (Corning, NY, USA). The cells were then maintained and aggregated as spheroids on an orbit-shaker (Thermo Fisher) at 100 RPM for over 7 days until analysis.

### 1.5 Glucose stimulated insulin secretion in stem cell-derived islets

SC-islets were hand-picked into transwells (CLS3414, Merck, Billercia, MA, USA) and placed in 24-well cell culture plates. The SC-islets were washed three times with Krebs buffer and equilibrated in Krebs buffer containing 2mM glucose for 60 min at 37 °C. The SC-islets were then sequentially incubated in Krebs buffer containing 2 mM and 20 mM glucose for 60 min each and the supernatant was collected. Last, the cells were incubated in Krebs buffer containing 20 mM glucose and 30 mM KCl for 30 min, the supernatant was collected. Cells were placed in a 37 °C humidified incubator with 5% CO_2_ for all incubations. Insulin in the supernatant was quantified with human insulin ELISA kit (Mercodia, Uppsala, Sweden). Prior to insulin determination with the LC-MS/MS method, the collected supernatants were spiked with a total of 5.0 pg/μL of bovine insulin and added 100% FA to a total of 0.5%.

### 1.6 Liquid chromatography instrumentation

The conventional LC-HESI-MS system was a modified Agilent 1100 series (Santa Clara, CA, USA) employing only shielded fused silica nanoViper™ (Thermo Fisher)) connectors in the entire system. Injection was achieved by coupling a 6-port-2-position valve, with a 50 μm inner diameter (id) x 550 mm nanoViper™ loop or a 20 μL nanoViper™ loop, between the output from the pump and in front of the column set-up. A 25 μL or 250 μL glass syringe was used for introduction of solutions and samples onto the loop. Introduction of method evaluation solutions onto the loop was done using a 150 μm id x 750 mm nanoViper™ coupled to a 250 μL glass syringe, where the solutions were filled via the nanoViper™. The column set-up consisted of an Accucore™ phenyl/hexyl guard column (2.1 mm id x 10 mm, 2.6 μm, 80 Å) attached with a Uniquard drop-in holder to the InfinityLab Poroshell EC-C18 separation column (2.1 mm id x 50 mm, 2.7 μm, 120 Å). For experiments with elevated column temperature, a 10 cm Butterfly portfolio heater coupled to a column heater controller, both from Phoenix S&T (Chester, PA, USA) was used.

### 1.7 Mobile phases and gradient settings for LC

The mobile phase (MP) reservoirs contained 0.1% FA in water added 1% DMSO (MP A) and 0.1% FA in ACN added 1% DMSO (MP B). A 150 μL/min solvent gradient was started at 1% B and linearly increased to 60% B in 8 min, further increased to 80% B at 8-12 min, quickly decreased to 40% B, and kept at 40% B for 4 min, before being further decreased to 1% B and kept at 1% B for 7 min. The gradient had a total runtime of 23 min.

### 1.8 Mass spectrometry instrumentation

A TSQ Quantiva triple quadrupole (QQQ) MS equipped with an H-ESI-II probe ionization source, both from Thermo Fisher, was used for all of the experiments. A syringe pump (model pump 11 elite) from Harvard Apparatus (Holliston, MA, USA) was used for direct injection on the MS by using a 250 μL glass syringe (Trajan Scientific and Medical, Ringwood, Australia) coupled to a 150 μm id x 750 mm nanoViper™. Initial experiments, prior to optimization with Box-Behnken, were attained with the MS operated in fullscanQ1 mode with the following default settings at 150 μL/min LC flow rate: The vaporizer temperature was set to 210 °C, and a spray voltage of 3.5 kV. Sheath gas was set at 27 arbitrary units (Arb), while auxiliary gas was set at 9 Arb, and sweep gas was not applied. The ion transfer tube temperature was kept at 325 °C.

#### 1.8.1 Optimization of peak areas in untargeted acquisition on the MS in fullscanQ1 mode with Box-Behnken experimental design

The H-ESI settings were optimized using a three-factor design with the MS operated in fullscanQ1 mode. A scan range from *m/z* 500 – 1500 was applied with 0.7 Q1 resolution. Box-Behnken was repeated twice to include six selected parameters related to the ionization source: Sheath gas, vaporizer temperature, and spray voltage were included in the first design, while the second design included sweep gas, auxiliary gas, and ion transfer tube temperature. Settings used for sweep gas, auxiliary gas and ion transfer tube temperature in the first set-up of Box-Behnken were the default settings used in the initial experiments, mentioned in **Section 2.8**. The three-factor designs were considered for the peak area of the most abundant precursor ion related to human insulin. The three-factor designs are described with the minimum and maximum value of each parameter in **Table 1**. The script written for Box-Behnken is available as an open source code using the code named “btjenesten” version 0.26 and “scikit-learn” version 1.0.2 found at *https://pypi.org/project/btjenesten/0.26/*.

**Table 1:** Description of parameters, including minimum and maximum value, used in two three-factor Box-Behnken designs for optimization of peak area of human insulin precursor.

| Parameter | Minimum value | Maximum value |
| --- | --- | --- |
| Sheath gas (Arb) | 20 | 36 |
| Vaporizer temperature (°C) | 170 | 250 |
| Spray voltage (kV) | 1.8 | 3.5 |
| Sweep gas (Arb) | 0 | 10 |
| Auxiliary gas | 5 | 13 |
| Ion transfer tube temperature (°C) | 275 | 375 |

#### 1.8.2 Targeted analysis with MS operated in selected reaction monitoring mode

The transitions, used in selected reaction monitoring (SRM), including collision energies, are listed in **Table 2**. The collision energies and radio frequency (RF) lens voltage were optimized using the compound optimization provided in Xcalibur. The collision gas (argon) pressure in q2 was 2.5 mTorr, the RF Lens was set at 210 V and the cycle time was set to 1 sec. The vaporizer temperature was set to 210 °C, and a spray voltage of 3.5 kV was applied to the H-ESI-II probe. Sheath gas was set at 20 Arb (approx. 2.71 L/min), while auxiliary gas was set at 9 Arb (approx. 7.49 L/min), and sweep gas was not applied. The ion transfer tube temperature was kept at 275 °C.

**Table 2:** SRM transitions used for human and bovine insulin (internal standard) including quantifier/qualifier status, precursor ion, product ion, and collision energy.

| **Compound** | **Identity** | **Precursor ion (*m/z*)** | **Product ion (*m/z*)** | **Collision energy (V)** |
| --- | --- | --- | --- | --- |
| **Human insulin** | Quantifier | 1162.5 | 226.1 | 41 |
|  | Qualifiers | 1162.5 | 345.1; 1159.0 | 42; 22 |
| **Bovine insulin** | Quantifier | 1147.8 | 226.2 | 41 |
| **(Internal standard)** | Qualifiers | 1147.8 | 315.2; 1144.5 | 42; 20 |

## 2 Results and discussion

### 2.1 Repeatable and stable fragmentation of intact insulin was attained on a QQQ mass spectrometer

Based on ELISA measurements, it was expected that insulin secretion from SC-islets would be in the lower pg/μL range, and that the developed LC-MS/MS method would need a ≤ 1 pg/μL detection limit. In addition, we aimed for developing a method which was independent of sample preparation, apart from adding internal standard to the samples. Therefore, effort was placed on optimizing the signal of the most abundant precursor of human insulin in the MS to achieve the lowest possible detection limit in biologically relevant matrices for characterization of SC-islet organoids (i.e. cell medium and Krebs buffer). Initial MS/MS settings were established: *m/z* 1162.5 (+5) →*m/z* 226.1, 345.0, 1159.2, 1358.0, see **Figure 1A**. These repeatable and stable MS/MS transitions, also commonly applied in insulin analysis [18, 19], allow for highly selective measurements, distinguishable from structural analogs e.g. insulin from animals, synthetic insulins, and proinsulin. See **SI-1** for more details.

**Figure 1:**
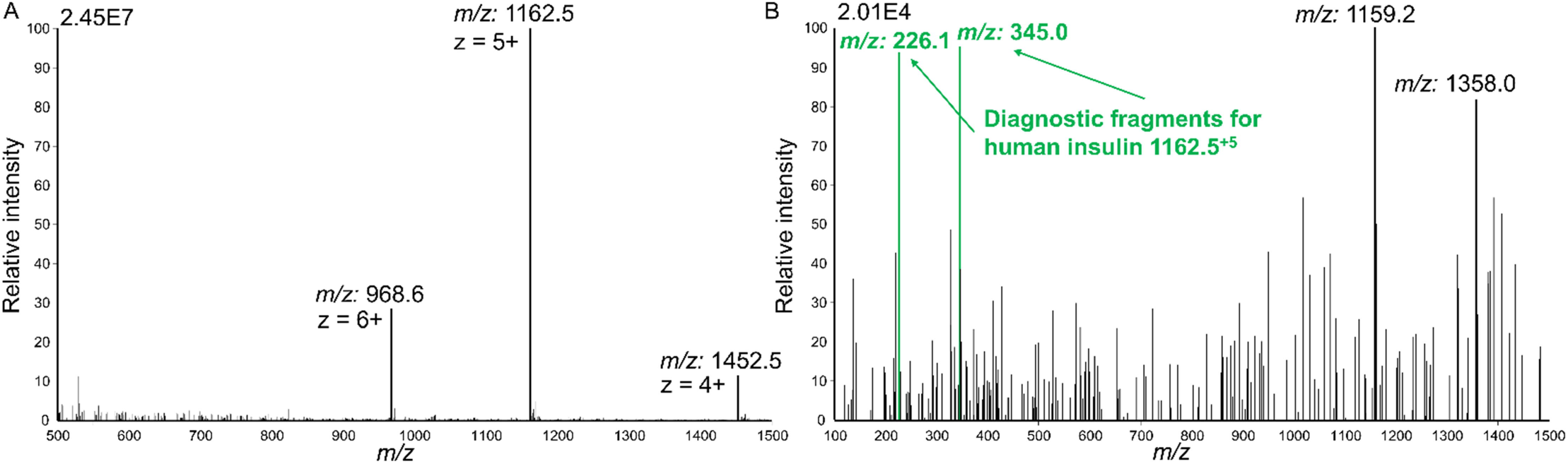
(**A**) Charge distribution of intact human insulin found in FullscanQ1 mode by continuously flow of a 1+1 mixture of acetonitrile and water with 10 ng/μL human insulin, (**B**) product ions fragmented from *m/z* 1162.5, the most abundant precursor of human insulin.

### 2.2 Non-defined adsorption of intact human insulin in autosampler and on glass syringe eliminated by shielded fused silica tubing

For initial attempts of direct injection of intact insulin in the previous **Section 2.1**, we examined a multitude of tubings (e.g. untreated fused silica capillary, polyetheretherketone (PEEK) tubing, fused silica/PEEK capillary from Agilent, and surface-treated PEEK shielded fused silica nanoViper™) to achieve a continuously stable signal of human insulin in the MS. A stable signal of human insulin was only obtained with the shielded nanoViper™, while insulin appeared to be retained on the other types of tubings due to non-defined adsorption (results not shown). Therefore, we replaced all available tubings in the LC-system with the shielded nanoViper™ tubing. However, some tubings (i.e. seat capillary, load capillary and a fused silica/PEEK capillary from Agilent) could not easily be replaced due to different formats not suitable for nanoViper™ fittings. The effect of the remaining unchangeable tubing was obvious when comparing autosampler injection with manual injection using a 6-port-2-position valve with an 1.08 μL nanoViper™ loop: for 125 pg/μL human insulin in an aqueous standard, poor signal was associated with the autosampler (**Figure 2A**), while a high intensity peak belonging to human insulin at *m/z* 1162.5 was attained with manual injection (**Figure 2B**). For injection of higher concentrations of insulin on the autosampler, the peak shape was drastically worse with an asymmetry factor of 2.4 (results not shown), while the peak obtained by manual injection had an asymmetry factor of 1.4 (**Figure 2B**). Similar chromatograms were obtained for bovine insulin (results not shown). In addition, we found that glass parts for e.g. syringes would also contribute to carry-over and should be avoided, visualized in **Figure 2C** (see **SI-2** for details). Although manual injections were performed here, contemporary autosamplers can be modified accordingly.

**Figure 2:**
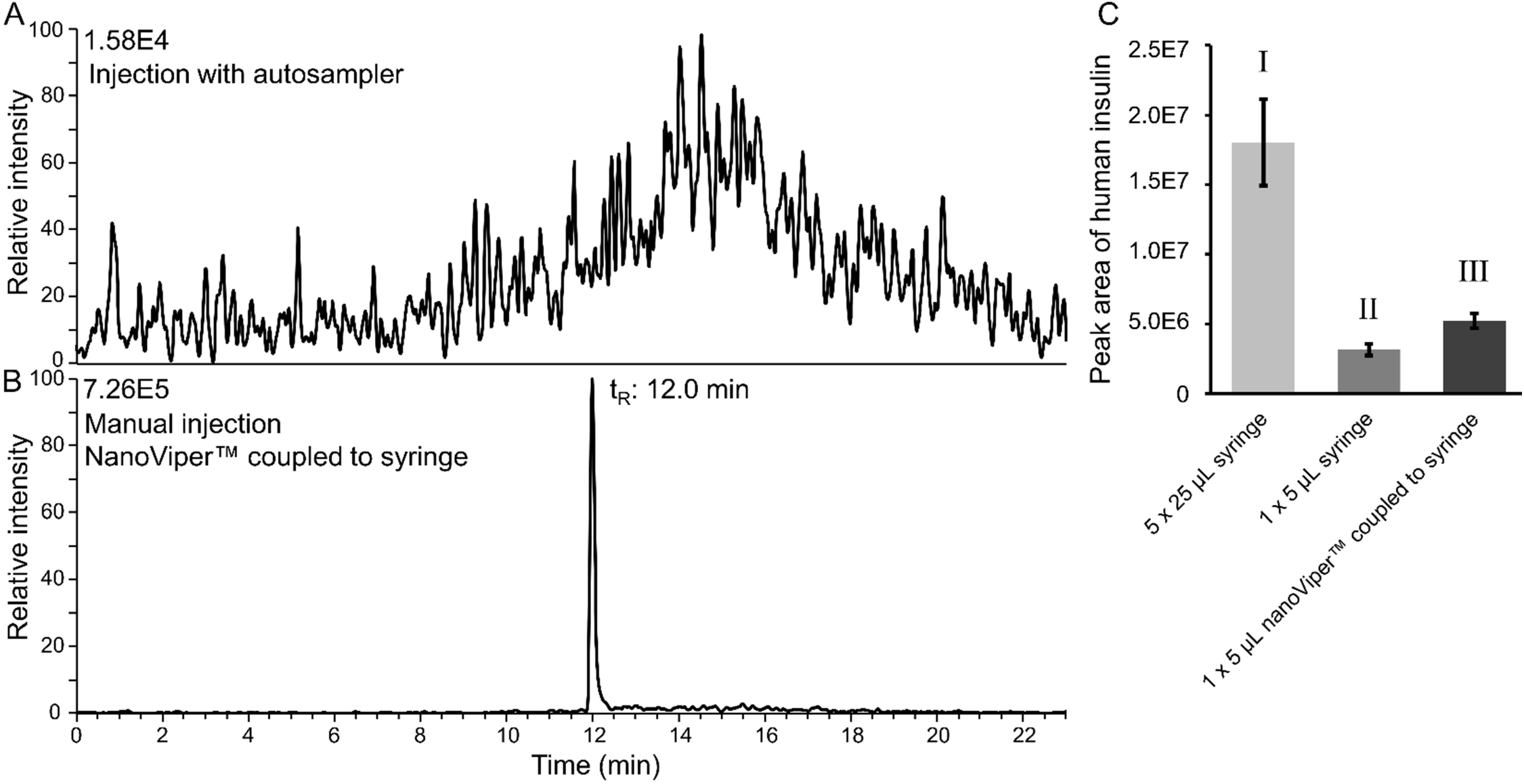
Extracted ion chromatogram of intact human insulin (*m/z* 1162.0-1163.0): (**A**) 1 μL injection of 125 pg/μL human insulin in 0.1% FA in water using autosampler, and (**B**) 1.08 μL injection of 125 pg/μL human insulin in 0.1% FA in water using a 6-port-2-position valve with 50 μm id x 550 mm nanoViper™ loop, and the solution was filled in the loop with a 25 μL glass syringe coupled to a 150 μm id x 750 mm nanoViper™. Solutions were examined with the MS operated in fullscanQ1 mode. (**C**) Comparison of peak areas of human insulin obtained with manual injection of 125 pg/μL insulin solution: (**I**) Shows that insulin accumulated on the syringe when 125 μL (5 × 25 μL) of solutions was used for wash and filling of the syringe and the loop prior to injection (n = 3), compared to (**II**) when the syringe was filled once with 25 μL and 5 μL applied over the 1.08 μL loop (1 × 5 μL, n = 3). (**III**) A compromise was coupling a 13.2 μL shielded fused silica nanoViper™ to the syringe, and filling 40 μL (2 × 20 μL) solution through the nanoViper™-syringe and applying 5 μL over the 1.08 μL loop (1 × 5 μL, n = 3).

### 2.3 Optimization of precursor peak areas with Box-Behnken led to lower detection limits

The peak area of precursor *m/z* 1162.5 of human insulin was optimized using Box-Behnken (BB) experimental design on the six selected variables in the H-ESI and MS-inlet settings. Column temperature was kept at ambient, as temperature did not affect peak shape or peak area, see **SI-3** for more details. Optimal settings found with BB for sheath gas (SG, Arb), vaporizer temperature (VT, °C), and spray voltage (SV, kV) were 36 Arb, 210 °C and 3.5 kV, see **Figure 3A**. However, by comparing the peak areas obtained with changing only the sheath gas levels from 36 Arb (**Figure 3B-I**) to 20 Arb (**Figure 3B-II**), an even higher peak area of human insulin was obtained. Optimized settings for sweep gas, auxiliary gas, and ion transfer tube temperature were found to be 0 Arb, 9 Arb and 275 °C (See **Figure 3C**). Comparing the initial settings to the optimized settings, sheath gas level was reduced from 27 Arb to 20 Arb and ion transfer tube temperature was decreased from 325 °C to 210 °C, which allowed a 3.2-fold sensitivity increase. See **SI-4** for a detailed description of the experiment design and evaluation.

**Figure 3:**
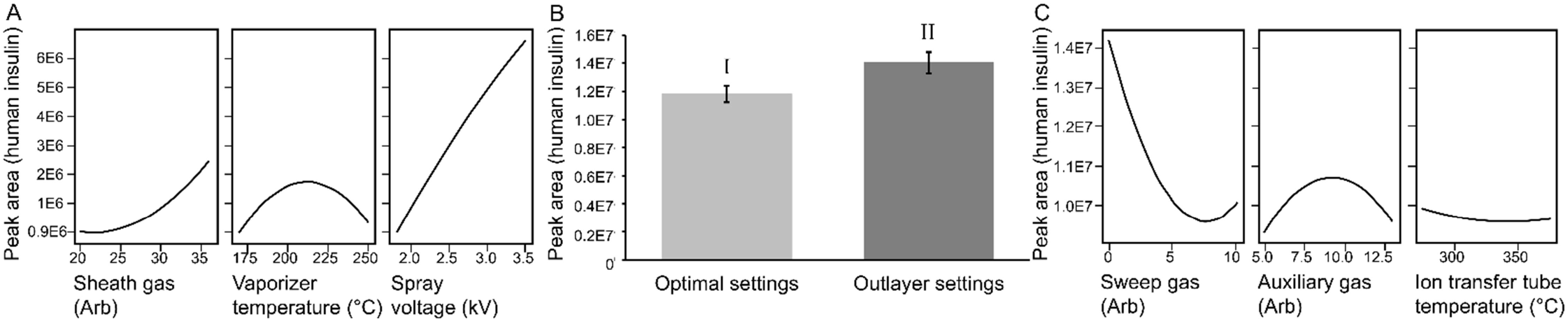
Optimization of peak area of human insulin *m/z* 1162.5 with Box-Behnken: (**A**) Main effects plot showing effect of sheath gas levels, vaporizer temperature and spray voltage, (**B**) comparison of peak areas obtained with (**I**) optimal settings [SG = 36, VT = 210, SV = 3.5] and (**II**) outlayer settings [SG = 20, VT = 210, SV = 3.5], and (**C**) main effects plot showing effect of sweep gas levels, auxiliary gas levels and ion transfer tube temperature.

### 2.4 Diagnostic fragment *m/z* 226 from human and bovine insulin established a calibration curve in cell medium for determining concentrations in quality controls with sufficient accuracy

The linearity, carry-over and the accuracy of the fully optimized SRM method with the following settings; SG = 20, VT = 210, SV = 3.5, SWG = 0, AUX = 9, ITT = 275, and collision gas pressure = 2.5 mTorr, were examined. Three transitions belonging to human insulin: *m/z* 1162.5 →*m/z* 226.1, *m/z* 345.0, or *m/z* 1159.2, and three transitions from bovine insulin: *m/z* 1147.8 →*m/z* 226.2, *m/z* 315.2, or *m/z* 1144.5, were monitored. A calibration curve was established in the range from 7.8 pg/μL to 250 pg/μL human insulin with 125 pg/μL bovine insulin as internal standard in cell medium for each possible combination of SRM transition from human and bovine insulin. Quality control (QC) samples with [10, 20, 50, 80, 100, 150, 200] pg/μL human insulin and 125 pg/μL bovine insulin were analyzed and the concentrations were determined using the established calibration curves. The calibration curves, which achieved the most accurate determination of the insulin concentrations in the QC’s, are shown with their respective SRM transitions for human and bovine insulin in **Figure 4A-C**. Only the QC standard with the lowest amount of human insulin (10 pg/μL) could not be determined with an accuracy less than 10% relative error. The combination of the transition *m/z* 1162.5 →*m/z* 226.1 for human insulin, and *m/z* 1147.8 →*m/z* 226.2 for bovine insulin (**Figure 4A**) provided on average an absolute relative error of 3% for the other six QC standards, and were therefore selected as the quantifiers. The four other remaining transitions *m/z* 1162.5 →*m/z* 345.0 or *m/z* 1159.2, and *m/z* 1147.8 →*m/z* 315.2, or *m/z* 1144.5, were used as qualifiers. Although an internal standard was employed in subsequent studies, it can be noted that also without the use of an IS, the precision and accuracy was excellent (see **SI-6**).

**Figure 4:**
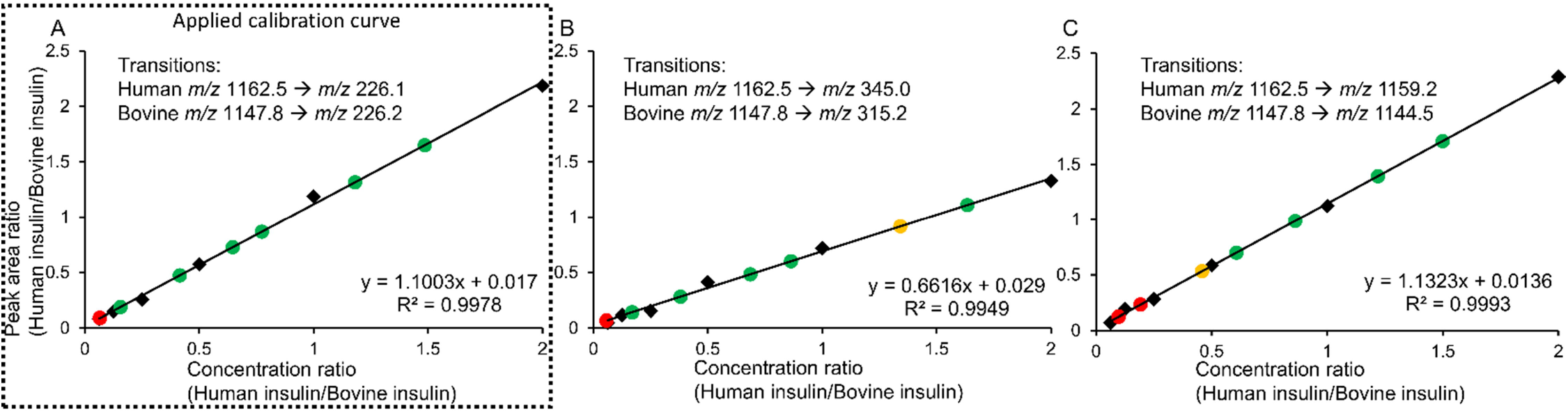
Calibration curve in 1.0% FA cell medium, obtained by SRM, from 7.8 pg/μL to 250 pg/μL human insulin with 125 pg/μL bovine insulin (IS) (points marked in black were used for calibration). Seven quality control standards at [10, 20, 50, 80, 100, 150, 200] pg/μL human insulin and 125 pg/μL bovine insulin are color-coded based on relative error (e_r_): green = e_r_ ≤ 10%), yellow = 10% < e_r_ < 15%, and red = e_r_ ≥ 15%. SRM calibration curves were established using the following transitions: (**A**) *m/z* 1162.5 → *m/z* 226.1 for human insulin, and *m/z* 1147.8 → *m/z* 226.2 for bovine insulin, (**B**) *m/z* 1162.5 → *m/z* 345.0 and *m/z* 1147.8 → *m/z* 315.2, and (**C**) *m/z* 1162.5 → *m/z* 1159.2 and *m/z* 1147.8 → *m/z* 1144.5.

The carry-over was examined after the injection of the calibration standard with the highest concentration of human insulin (250 pg/μL). For all of the six transitions applied as quantifier or qualifier, carry-over in the following blank was ≤ 3%.

### 2.5 Detection limit below 1 pg/μL was reached in Krebs buffer by increasing the injection volume

To reach the target detection limit below 1 pg/μL, an increase in the injection volume was examined. The increase in injection volume was easily achieved by exchanging the 1.08 μL loop with a 20 μL loop and using only a syringe for injection (see **SI-5** for discussion concerning use of only a syringe for injection). The increase in injection volume caused a collapse of the chromatography for standards in cell medium. With 1.08 μL injection of a solution consisting of 125 pg/μL human and 125 pg/μL bovine insulin, insulins were eluted at 11.2 min and were separated from the other compounds in the cell medium, which were eluted after 11.6 min, see **Figure 5A**. When the injection volume was increased to 20 μL of a solution consisting of 6.25 pg/μL human and 6.25 pg/μL bovine insulin, no peaks corresponding to human or bovine insulin was distinguishable from the other compounds present in cell medium, as shown in **Figure 5B**.

**Figure 5:**
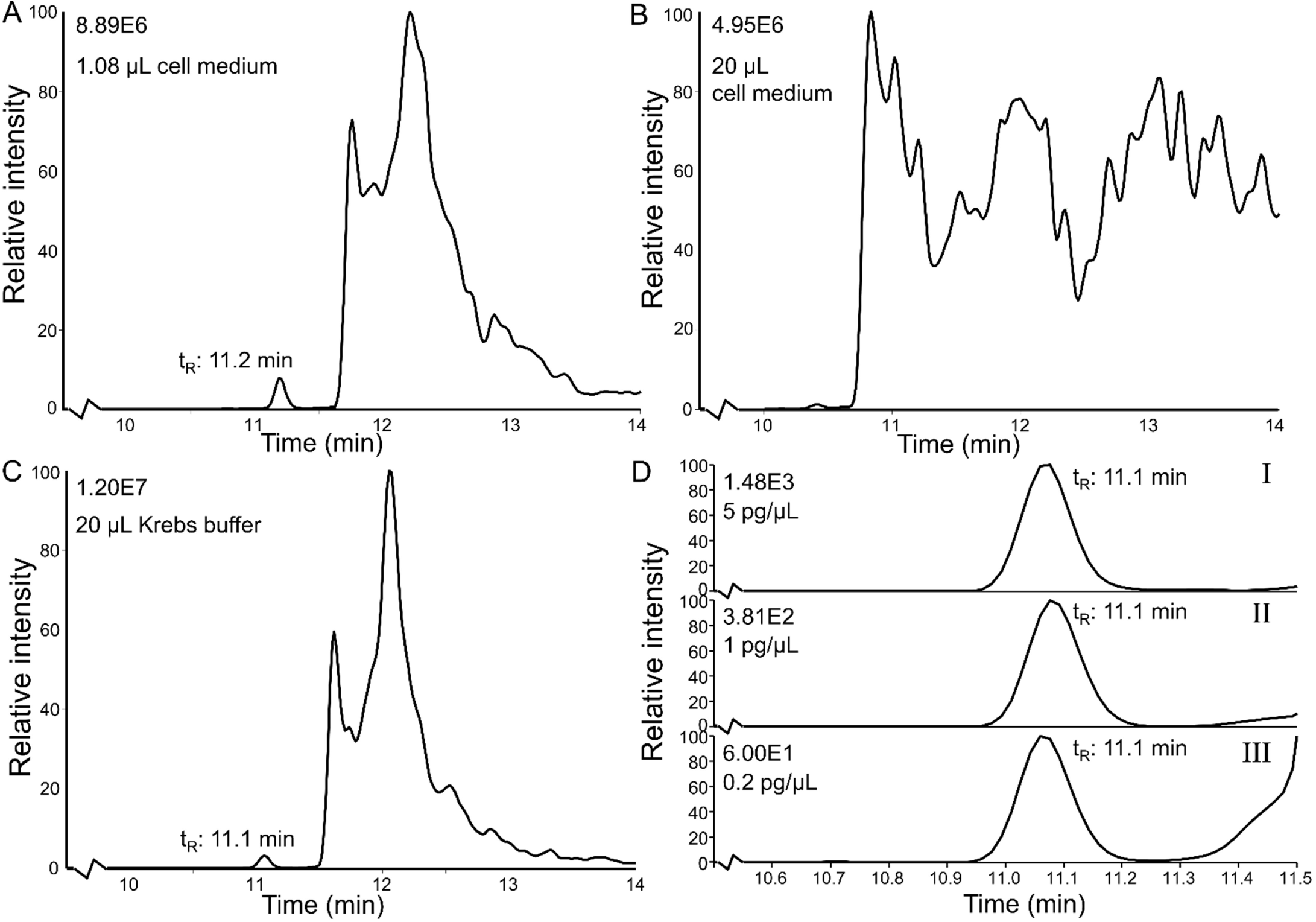
Effect of injection volume and sample matrix on the separation was examined in FullscanQ1 mode. Extracted ion chromatogram (*m/z* 1162-1163) showing: (**A**) Human insulin detected at 11.2 min after 1.08 μL injection of 125 pg/μL human insulin in 1% FA in cell medium, (**B**) human insulin not detected in 20 μL injection of 6.25 pg/μL human insulin in 1% FA in cell medium, and (**C**) human insulin detected at 11.1 min after 20 μL injection of 5 pg/μL human insulin in 0.5% FA in Krebs buffer. (**D**) Peaks obtained in SRM mode of human insulin in a dilution series in 0.5% Krebs buffer with the quantifier transition *m/z* 1162.5 → *m/z* 226.1: (**I**) 5 pg/μL human insulin, (**II**) 1 pg/μL human insulin, and (**III**) 0.2 pg/μL human insulin.

Besides cell medium, Krebs buffer is another biologically relevant matrix for assessing SC-islets. When 20 μL of Krebs buffer with 5 pg/μL human and 5 pg/μL bovine insulin was injected, human and bovine insulin were eluted at 11.1 min and were separated from other compounds in the buffer and residue of cell medium on the column, shown in **Figure 5C**. The detection limit was successfully lowered, shown here by detection of human insulin in a dilution series in Krebs buffer at the following concentrations: 5 pg/μL (**Figure 5D-I**), 1 pg/μL (**Figure 5D-II**), and 0.2 pg/μL (**Figure 5D-III**), with the MS operated in SRM mode. Results shown in **Figure 5D** used the quantifier SRM transition (*m/z* 1162.5 →*m/z* 226.1), but detection was also successful with the other selected qualifier transitions (*m/z* 1162.5 →*m/z* 345.0, or *m/z* 1159.2).

Thus by increasing the injection volume to 20 μL, a detection limit of ≤ 0.2 pg/μL (34.4 pM) was achieved in standards prepared in Krebs buffer with 0.5% FA. For standards prepared in cell medium with 1.0% FA, the increased injection volume compromised the separation. Cell medium and Krebs buffer are common matrices used in characterization of human islets and islets organoids, and we found that with our LC-MS method the highest sensitivity for insulin analysis were achieved in Krebs buffer.

### 2.6 Optimized LC-MS/MS method determined insulin secretion in stem cell-derived islets

To determine insulin secretion from SC-islets (**Figure 6A-B**, and see more details in **SI-7**) exposed to three different glucose environments, a calibration curve was established from 0.2 pg/μL to 10 pg/μL human insulin with 5.0 pg/μL bovine insulin as internal standard, see **Figure 6C**. The accuracy of the calibration and potential drift in the system during a long sample sequence was monitored by running a quality control standard with 2.0 pg/μL human insulin (with 5.0 pg/μL bovine insulin as internal standard) as the first injection following the calibration standards, in the middle of the sample sequence and as the last injection. The quality controls were individually determined with a relative error of -2%, -9% and 2%, and the average concentration was found to be 1.9 pg/μL with an RSD of 6%. The quality controls indicated that the LC-MS/MS method provide sufficient accuracy in determining insulin concentrations by a single injection of a sample and that there was no drift in the system in the course of a relatively long sequence (37 injections).

**Figure 6:**
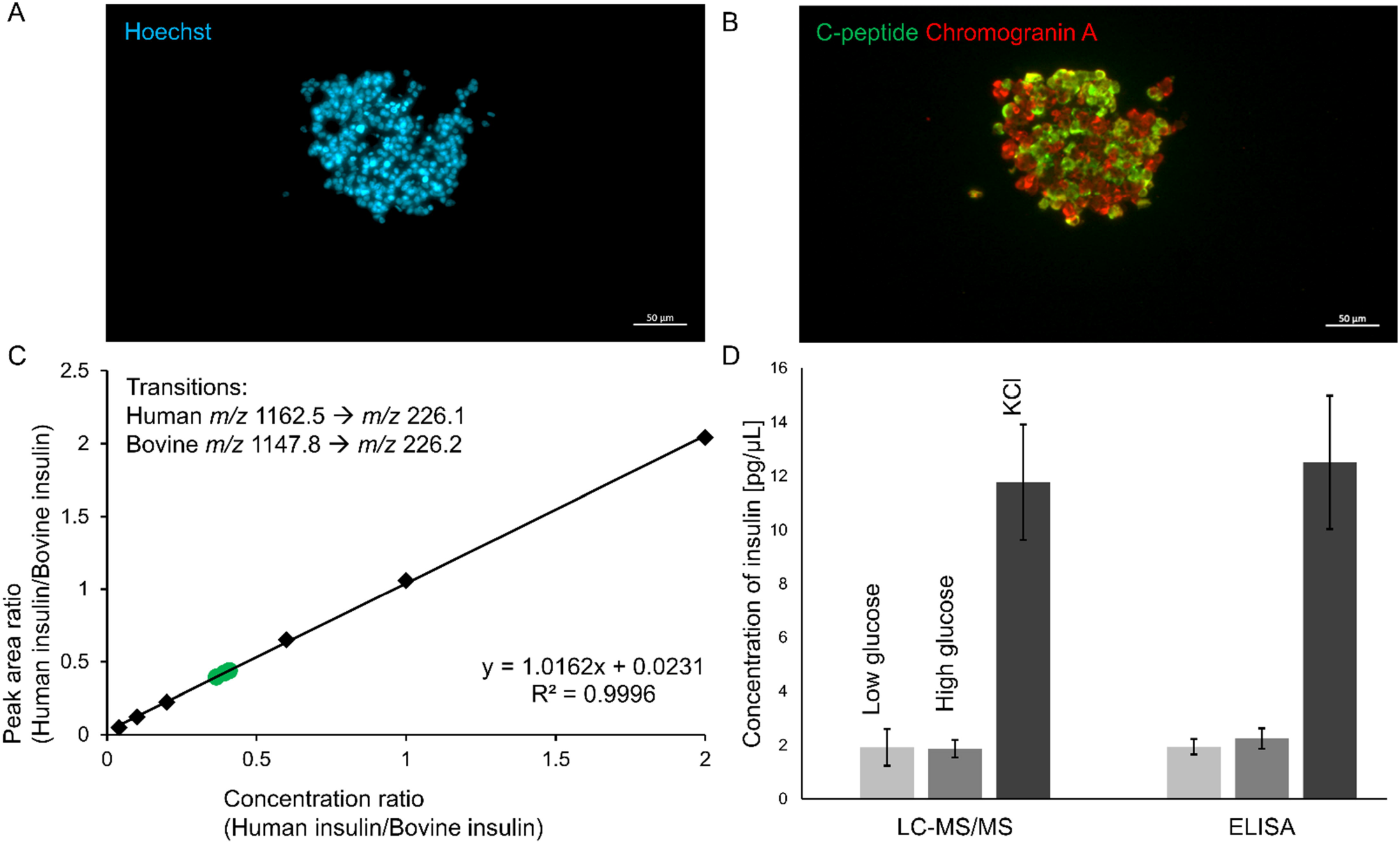
Representative immunostaining images of SC-islets: (**A**) Nuclei DNA stained with Hoechst (blue), and (**B**) endocrine secretory vesicles stained with chromogranin A (red) and production of insulin is represented by C-peptide (green). (**C**) Calibration curve in 0.5% FA Krebs buffer, obtained by SRM, from 0.2 pg/μL to 10 pg/μL human insulin with 5 pg/μL bovine insulin (IS) (points marked in black), and three quality control standards (marked green as relative error was less than 10%) at 2 pg/μL human insulin with 5 pg/μL bovine insulin (IS). (**D**) Insulin secretion determined by LC-MS/MS and ELISA from SC-islets exposed to low glucose (2 mM), high glucose (20 mM) and KCl (20 mM glucose with 30 mM KCl).

Analyses by the LC-MS/MS method showed that SC-islets exposed to 2 mM glucose (low glucose) secreted 1.9 pg/μL of human insulin (n = 4, RSD = 36%). Contradictory, in 20 mM glucose (high glucose), 1.9 pg/μL of human insulin (n = 4, RSD = 18%) was also secreted by the SC-islets. However, when the SC-islets were exposed to 20 mM glucose combined with 30 mM KCl, a total of 12 pg/μL of human insulin (n = 4, RSD = 18%) was secreted (it should be noted that the concentration of insulin in the KCl samples were outside of the calibration range). The values obtained by the LC-MS/MS method was compared to that of an established ELISA method, which determined human insulin secretion to be 2.0 pg/μL (n = 4, RSD = 15%) in low glucose, 2.3 pg/μL (n = 4, RSD = 17%) in high glucose, and 13 pg/μL (n = 4, RSD = 20%) in KCl environment. An independent two sample t-test, at 95% confidence, showed that the average insulin concentration found for the three different exposures determined with ELISA and LC-MS/MS were not significantly different, and this is visualized in **Figure 6D**. For quality SC-islets, it was expected that the insulin secretion in response to high glucose should be more than 1.5 times higher than the insulin secretion in response to low glucose. The SC-islets used in these experiments did not show this difference in insulin secretion based on glucose environment. However, it should be noted that the SC-islets used in these experiments were examined in the first days after completion of the differential protocol, and we would expect SC-islets that have matured for a couple more days to display a difference in insulin secretion profile depending on glucose concentration. The experiments with KCl show however, that the SC-islets are producing larger amount of insulin, and are able to control how much is secreted pending on exposure. The LC-MS/MS method shows sufficiently low detection limit as we are able to detection insulin secretion from not functionally mature SC-islets.

## 3 Brief comparison to previously published LC-MS methods for determination of human insulin

In a balanced salt solution, equivalent to the Krebs buffer applied in this study, Donohue *et al*. [16] developed a conventional RPLC-MS/MS method for human insulin with a precision of RSD = 13-16% and a detection limit of 0.5 nM (2.9 pg/μL). Another multi-element conventional RPLC-MS/MS method achieved simultaneous detection of human insulin, synthetic analogous, and C-peptide from blood/plasma samples by combining protein precipitation and a mixed-mode cation-exchange solid-phase extraction sample preparation. The method was fully validated with a LOD of 0.2 pg/μL, LOQ of 0.6 pg/μL, 78-128% accuracy, and < 21% precision [18].

The method presented in this study, tailored for islet organoid samples, has negligible carry-over, an LOD of ≤ 0.2 pg/μL, ≤ ±10% accuracy, and < 10% precision for human insulin, which is in the same range as achieved in the previously mentioned methods.

## 4 Conclusions

We have developed an RPLC-MS/MS method tailored for measuring insulin secreted from organoid islets. The method is highly sensitive and selective, with excellent quantitative properties. Conventional LC columns were compatible with low μL-scale samples, and next steps will be direct coupling of the system with pancreas-on-a-chip systems [8]. A limitation is that the method is sensitive to the type of medium applied; additional fine-tuning can eventually be performed, e.g. focusing on LC column selectivity to avoid medium specific matrix effects. Possible expansions include adding additional analytes to the method, for example cytokines and other islets-related peptides (e.g. somatostatin, glucagon, and C-peptide).

## Acknowledgements

Financial support was obtained from the Research Council of Norway through its Centres of Excellence funding scheme, project number 262613 and partly from UiO:Life Science. S.R.W. is a member of the National Network of Advanced Proteomics Infrastructure (NAPI), which is funded by the Research Council of Norway INFRASTRUKTUR-program (project number: 295910).

## Conflict of interest statement

The authors declare no conflict of interest.

## Notes

### Competing Interest Statement

The authors have declared no competing interest.

