## Supporting Information for "Determination of insulin secretion from stem cell-derived islet organoids with liquid chromatography-tandem mass spectrometry"

***Supplementary Material for*: Determination of insulin secretion from stem cell-derived islets with liquid chromatography-tandem mass spectrometry**

Christine Olsen^a,b^, Chencheng Wang^b,c^, Shadab Abadpour^b,c^, Elsa Lundanes^a^, Audun Skau Hansen^a^, Frøydis Sved Skottvoll^d^, Hanne Scholz^b,c^, and Steven Ray Wilson^a,b*^

^a^University of Oslo, Department of Chemistry, Blindern, Oslo, Norway

^b^Hybrid Technology Hub-Centre of Excellence, Institute of Basic Medical Sciences, Faculty of Medicine, University of Oslo, Oslo, Norway

^c^Department of Transplant Medicine and Institute for Surgical Research, Oslo University Hospital, Oslo, Norway

^d^Department of Smart Sensors and Microsystems, SINTEF Digital, Oslo, Norway

### 1 Discussion concerning selection of collision gas pressure for stable fragmentation of intact insulin

Assessment of an aqueous standard containing 10 ng/µL human insulin by direct injection on a QQQ MS operated in fullscanQ1 mode showed that intact insulin (*i.e.* no changes to the structure of the peptide from its natural form) was present with expected charge states: *m/z* 968.6 (z = +6), *m/z* 1162.5 (z = +5) and *m/z* 1452.5 (z = +4), see **Figure S1A**. The fragmentation of the most abundant ion *m/z* 1162.5 was examined at various collision gas pressures from 1 mTorr to 4 mTorr. Commonly used diagnostic fragments in targeted insulin determination (*i.e.* *m/z* 226.1 and *m/z* 345.0) and larger fragments (*m/z* 1159.2 and *m/z* 1358.0, see **Figure S1B**) was found with increasing intensity when applying increasing collision gas pressures from 1.0 mTorr to 3.0 mTorr, see **Figure S1C** [1, 2]. At gas pressures of 2.5 mTorr and 3.0 m Torr, the signal intensity of the smaller fragments were in equal range, but with a larger intensity variance at 3.0 mTorr. The larger fragment *m/z* 1159.2 was present at sufficient intensities at both 2.5 mTorr and 3.0 mTorr gas pressures. At higher gas pressures over 3.5 mTorr, more of the precursor ion remained intact and the larger product ions were present at a higher intensity, however, the smaller fragment ions *m/z* 226.1 and *m/z* 345.0 were not present in all of the product scans. Therefore, 2.5 mTorr was selected as the collision gas pressure in the targeted experiments.

Bovine insulin has three amino acids which are different compared to that of human insulin and was selected as the internal standard in the method due to its availability and being an inexpensive alternative to a deuterated compound. A similar evaluation of bovine insulin by direct injection showed a highly abundant and stable precursor at *m/z* 1147.8 (z = +5) and high-intensity product fragments at *m/z* 226.2, *m/z* 315.2, and *m/z* 1144.5 at 2.5 mTorr collision gas pressure.

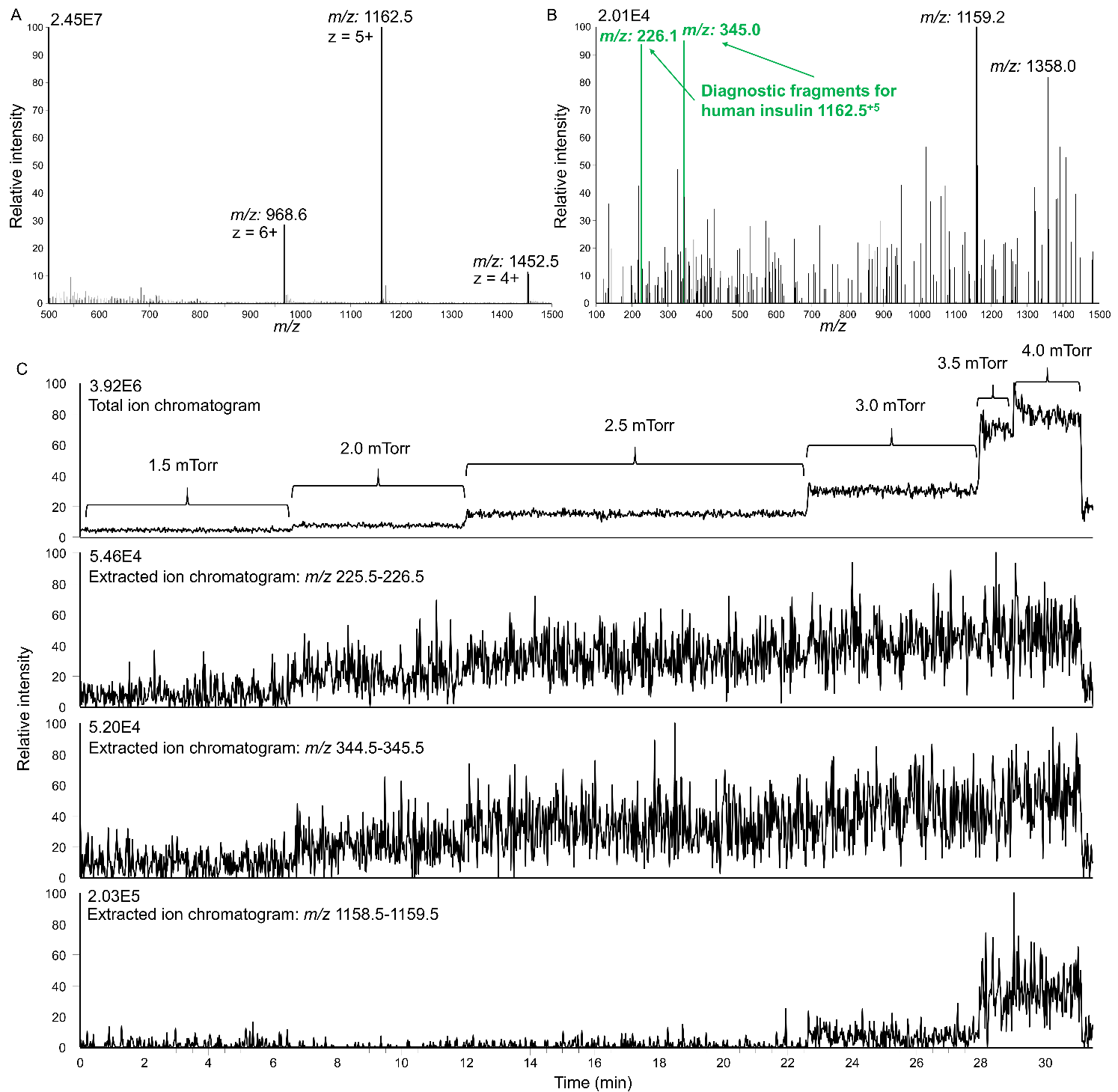

**Figure S1:** (**A**) Charge distribution of intact human insulin found in FullscanQ1 mode, (**B**) product ions fragmented from *m/z* 1162.5, the most abundant precursor of human insulin, and (**C**) evaluation of the stability and signal intensity of three selected product ions (i.e. *m/z* 226.1, *m/z* 345.0, and *m/z* 1159.2) from *m/z* 1162.5 at different collision gas pressures (1.5 mTorr – 4.0 mTorr).

### 2 Non-defined adsorption of intact human insulin on glass syringe eliminated by shielded fused silica tubing

The glass syringe used for manual injection was another critical point to assess for non-defined adsorption of insulins. Initially, insulin solutions were introduced directly onto the loop with a 25 µL glass syringe. When the glass syringe and the loop was washed with 25 µL of the insulin solution five times prior to loading and injection, the average peak area of human insulin was 1.8E7 with a relative standard deviation (RSD) of 17% (n = 3, see **Figure S2C-I**). The peak areas of human insulin also steadily increased for each subsequent injection: 1.5E7, 1.8E7 and 2.1E7. By reducing the amount of insulin solution filled on the syringe, syringe filled only once with 25 µL and directly fill 5 µL onto the loop for injection, the average peak area of human insulin was 3.2E6 with a RSD of 13% (n = 6, see **Figure S2C-II**). Seeing as the peak areas varied 83% between the two injection procedures, the glass syringe and the loop was examined for insulin accumulation by assessing the signal in a blank injection following a regular insulin injection. The carry-over signal of human insulin was ≥ 9% when only the loop had been cleaned with a 50/50 MeOH/H_2_O solution and 0.1% FA in water, while the carry-over signal was ≤ 2% when only the glass syringe was cleaned with 50/50 MeOH/H_2_O solution prior to filling with blank solution 0.1% FA in water. The carry-over indicated that insulin displayed non-defined adsorption towards the surface of the glass syringe, but not on the shielded nanoViper™ loop.

To avoid insulin accumulation on the glass syringe during injection, the syringe was coupled to a 150 µm id x 750 mm shielded nanoViper™ (volume of tubing = 13.2 µL). When the insulin solution was filled into the glass syringe through the nanoViper™ (two times 20 µL), and 5 µL of the volume in the tubing was filled onto the loop for injection, the average peak area of human insulin was 5.2E6 with a RSD of 10% (n = 6, **Figure S2C-III**). An independent two sample t-test, at 95% confidence, showed that the average peak area found for human insulin by injection with and without a nanoViper™ attached to the syringe were significantly different. In addition, the average peak area of human insulin obtained with injection using a syringe coupled to a nanoViper™ was between the average peak areas achieved with the previously examined injection procedures. Hence, it was even more probable that insulin had non-defined adsorption on the glass syringe, which accumulated during multiple fillings of the syringe giving a higher peak area, and when the syringe was filled only once led to a reduced peak area.

Injection technique and selection of tubing was of great importance in achieving LC-MS determination of intact insulin in low pg/µL range. A fully modified system, where insulin only came in contact with shielded fused silica nanoViper™, offered a repeatable chromatographic performance based on peak area examination. Utilizing a nanoViper™ coupled to the syringe eliminated the effect of non-defined adsorption on the syringe, and possible carry-over from the glass syringe was easily removed by washing with a 1+1 MeOH/water (v/v) solution.

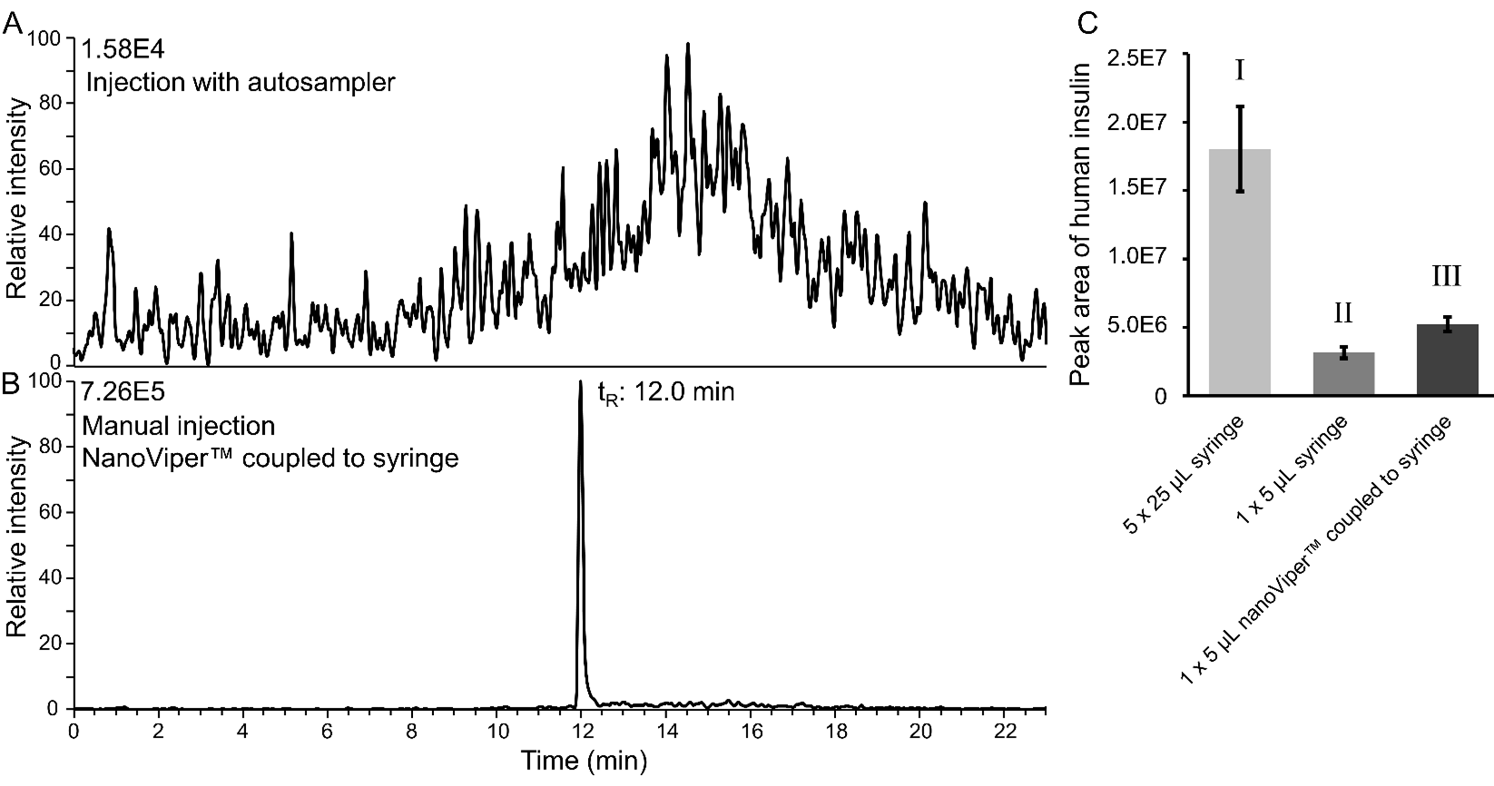

Figure S2: Extracted ion chromatogram of intact human insulin (*m/z* 1162.0-1163.0): (A) 1 µL injection of 125 pg/µL human insulin in 0.1% FA in water using autosampler, and (B) 1.08 µL injection of 125 pg/µL human insulin in 0.1% FA in water using a 6-port-2-position valve with 50 µm id x 550 mm nanoViper™ loop, and the solution was filled in the loop with a 25 µL glass syringe coupled to a 150 µm id x 750 mm nanoViper™. Solutions were examined with the MS operated in fullscanQ1 mode. (C) Comparison of peak areas of human insulin obtained with manual injection of 125 pg/µL insulin solution: (I) Syringe and loop filled five times with 25 µL, (II) syringe filled once with 25 µL and 5 µL applied on loop, or (III) 40 µL solution filled through a 13.2 µL nanoViper™ coupled to the syringe, and 5 µL were applied on the loop.

### 3 Discussion concerning effect of column temperature on peak area or peak shape

To examine if insulin peak shape/areas could be affected by the column temperature, the guard cartridge and separation column was placed in a column heater. Prior to injections, heating was applied for one hour to avoid a temperature gradient in the cross section of the columns. By injection of 1.08 µL of 125 pg/µL of human insulin and 125 pg/µL of bovine insulin dissolved in cell medium, no difference in peak area was found between 20 °C, 30 °C and 40 °C, as the average peak areas of human insulin was 4.4E6 (n = 3), 4.3E6 (n = 3) and 4.3E6 (n = 2), respectively. There was no difference found for bovine insulin either as the average peak areas of *m/z* 1147.8 was 4.2E6 (20 °C, n = 2), 4.1E6 (30 °C, n = 3) and 4.5E6 (40 °C, n = 2). Concerning the peak shape, no obvious difference was seen in the asymmetry factor as the factor was calculated to be 1.25 (n = 3) at 20 °C, 1.23 (n = 3) at 30 °C and 1.14 (n = 2) at 40 °C for human insulin.

### 4 Expanded discussion concerning optimization of precursor peak areas with Box-Behnken

The peak area of precursor *m/z* 1162.5 of human insulin was optimized using Box-Behnken (BB) experimental design on the six selected variables in the H-ESI and MS-inlet settings.

In the first set-up of a three factor BB-design: Sheath gas (SG, Arb) [20, 28, 36], vaporizer temperature (VT, °C) [170, 210, 250], and spray voltage (SV, kV) [1.8, 2.65, 3.5] was evaluated for the highest peak area of *m/z* 1162.5 in water standards. The optimized settings were found to be [SG = 36, VT = 210, SV = 3.5], as shown in main effects plot in **Figure S3A**. The optimized settings were included as one of the experiments in the BB-design and produced a peak with an area of 5.5E6. However, one of the experiments [SG = 20, VT = 210, SV = 3.5] (referred to as outlayer settings) achieved a peak area of 9E6, which was 39% larger than the peak area found by the settings deemed optimal by BB. In addition, as is common in BB, three experiments were run at the same settings, [SG = 28, VT = 210, SV = 2.65], which had an average peak area of 4E6 with a large variation, RSD = 34%. There was also a significant signal found in the blank injection following the 15 injections of 125 pg/µL human and 125 pg/µL bovine insulin, where the signal was equal to 7% carry-over of human insulin and 8% carry-over of bovine insulin. Therefore, to select between the optimized settings and the outlayer settings, three additional replicates of each experiments were compared after allowing the selected settings to run for one hour to reach a more stable condition, and a blank injection was run between each injection of insulin solutions. The optimized settings from Box-Behnken resulted in a peak area of *m/z* 1162.5 = 1.18e7 (RSD = 5%, n = 3, **Figure S3B-I**), while the outlayer settings showed a peak area of *m/z* 1162.5 = 1.40e7 (RSD = 5%, n =3, **Figure S3B-II**). The carry-over after a single injection was between 1-3% for both human and bovine insulin (n = 3), with no difference between the optimal and the outlayer settings. An independent two sample t-test, at 95% confidence, showed that the average peak areas of the two settings were significantly different, and therefore the outlayer settings were selected to be applied in further optimization.

In the second set-up of a three factor BB design: Sweep gas (SWG, Arb) [0, 5, 10], auxiliary gas (AUX, Arb) [5, 9, 13], and ion transfer tube temperature (ITT, °C) [275, 325, 375] were examined in standards prepared in cell medium. The optimized H-ESI settings for *m/z* 1162.5 were found to be [SWG = 0, AUX = 9, ITT = 275], as shown in main effects plot in **Figure S3C**, and had a peak area of *m/z* 1162.5 = 1.7E7. The three experiments, which were examined at the same settings; [SWG = 5, AUX = 9, ITT = 325] gave an average peak area of 1.1E7 with an RSD = 22% in this case. Another important feature shown for the method with the second BB-design, was that the blank injection following 15 injections of 125 pg/µL human and 125 pg/µL bovine insulin in cell medium had a signal of human insulin equal to 1% carry-over and there was no distinguishable peak corresponding to bovine insulin, indicating that the carry-over was distinctly reduced in cell medium standard compared to water standards.

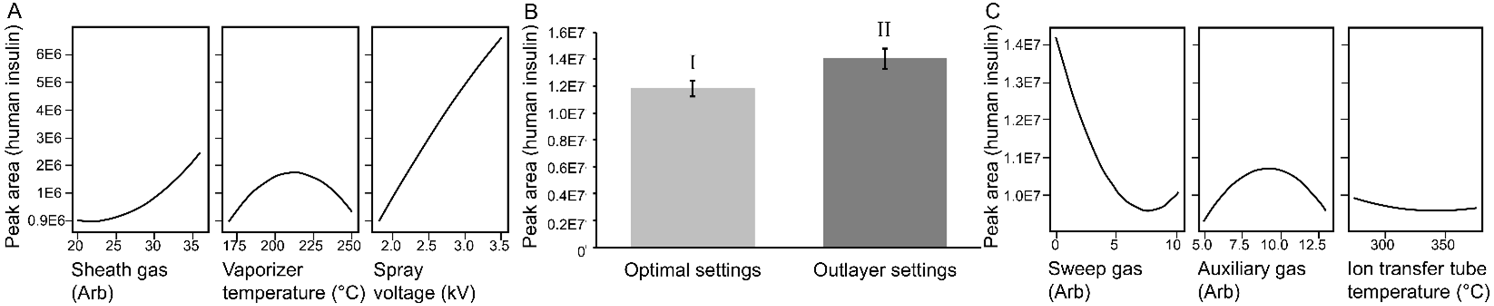

Figure S3: Optimization of peak area of human insulin *m/z* 1162.5 with Box-Behnken.: (A) Main effects plot showing effect of sheath gas levels, vaporizer temperature and spray voltage, (B) comparison of peak areas obtained with (I) optimal settings [SG = 36, VT = 210, SV = 3.5] and (II) outlayer settings [SG = 20, VT = 210, SV = 3.5], and (C) main effects plot showing effect of sweep gas levels, auxiliary gas levels and ion transfer tube temperature.

### 5 Discussion concerning manual injection technique based on peak areas obtained in SRM mode

The reduced carry-over of human and bovine insulin when working with standards prepared in cell medium compared to water standards, as described in previous **Section SI-2 and SI-4**, indicated that compounds present in the cell medium reduced the non-defined adsorption of insulins. Bovine serum albumin (BSA) was a possible candidate to examine, as BSA has been described as a sacrificial protein to block adsorption of proteins and peptides [3]. To examine if BSA could block non-defined adsorption, water standards with 125 pg/µL human and 125 pg/µL bovine insulin was prepared with and without 5 ng/µL BSA. The two solutions were then injected using only a glass syringe and a glass syringe coupled to a nanoViper™, a repeat of the same experiments used to establish that the glass syringe showed non-defined adsorption of insulin in water standards in **Section SI-2**. The experiments was done using SRM, as it was assumed SRM had a higher sensitivity, than a FullscanQ1 method, to examine potential carry-over, and SRM was the intended method to be applied in determination of insulin in SC-islets. Water standards without BSA injected with a nanoViper™ coupled to a glass syringe resulted in an average peak area for human insulin of 1.4E4 (n = 3, RSD = 10%), and injection using only a glass syringe gave a similar average peak area and variance (1.4E4 and RSD = 13%). The carry-over in a blank injection after three injections was in both cases less than 0.5%. When BSA was added to the water standard and injected with a nanoViper™ coupled to a glass syringe the average peak area for human insulin was 1.5E4 (n = 3, RSD = 7%), while injection using only a glass syringe gave a similar average peak area and variance (1.4E4 and RSD = 2%). The carry-over in a blank injection after three injection of BSA water standard was also less than 0.5% independent of injection technique. By two-way analysis of variance (ANOVA), there was no significant difference in the obtained average peak areas with or without BSA or depending on the injection technique. From the information gained when comparing the injection techniques and the presence of BSA, it could not be confirmed whether or not BSA reduced non-defined adsorption of human insulin.

The injection technique with the nanoViper™ coupled to the glass syringe is much more laborious and more exposed to errors in the execution than injection with only a glass syringe. In addition, more sample is required when the nanoViper™-syringe technique is applied and frothing was an issue with the cell medium standards. Therefore, injection of cell medium standards were examined with both injection techniques as well to see if the peak areas where significantly different or not with the SRM method. Cell medium standards injected with nanoViper™-syringe technique had an average peak area of 1.56E4 (n = 3, RSD = 2%) for human insulin, while injection with only the syringe had an average peak area of 1.8E4 (n = 3, RSD = 8%). An independent two sample t-test, at 95% confidence, showed that the average peak area of human insulin found for the two different injection techniques were not significantly different for cell medium standards. The standard deviations of the two injection techniques were also not significantly different. Based on these findings, only a glass syringe was used for injection of samples and standards in cell medium.

### 6 LC-MS/MS method sufficiently robust to maintain accuracy and precision without internal standard

The LC-MS/MS method robustness was evaluated by removing the internal standard and examining whether the method provided sufficient accuracy and precision without the internal standard. The three replicates of the 2.0 pg/µL human insulin quality control standard were individually determined with a relative error of -7%, -11% and -11%, and the average concentration was found to be 1.9 pg/µL with an RSD of 11%, results visualized in **Figure S4**. The quality controls indicates that the accuracy and precision of the LC-MS/MS method was not compromised by removing the internal standard. Without internal standard, the LC-MS/MS determined human insulin secretion to be 1.8 pg/µL (n = 4, RSD = 37%) in low glucose, 1.8 pg/µL (n = 4, RSD = 21%) in high glucose, and 11.3 pg/µL (n = 4, RSD = 14%) in KCl environment. An independent two sample t-test, at 95% confidence, showed that the average insulin concentration found for the three different exposures determined by LC-MS/MS with and without internal standard were not significantly different.

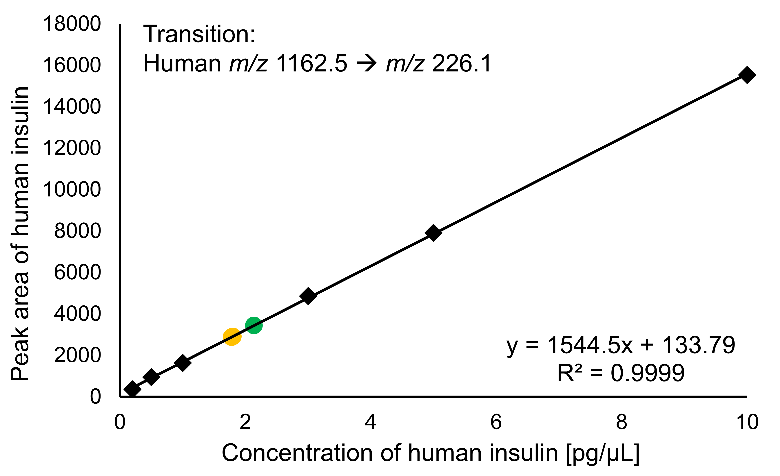

Figure S4: Calibration curve in 0.5% FA Krebs buffer, obtained by SRM, from 0.2 pg/µL to 10 pg/µL human insulin without internal standard(points marked in black), and three quality control standards at 2 pg/µL human insulin. Two quality controls are marked in yellow as the relative error was -10% and -11%, and the third QC is marked in green as the relative error was 7%.

### 7 Characterization of stem cell-derived islets

The SC-islets were generated from human pluripotent cell line H1 and maintained in suspension culture condition over 7 days until analysis. As shown in the bright field image, SC-islets were uniform in shape, with a diameter of around 100 µm at day 7 of aggregation (**Figure S5A**). Before being used for glucose-stimulated insulin secretion assay, SC-islets were further characterized with flow cytometry and immunofluorescence staining. Primary antibodies including human C-peptide (Developmental studies Hybridoma Bank, University of Iowa, IA, USA) and chromogranin A (CHGA, Novus Biologicals, Centennial, CO, USA) represent insulin-producing cells and pan-endocrine cells in the organoids respectively [4]. Representative flow cytometry quantification shows that > 96% of cells are endocrine cells (Q2 and Q3, **Figure S5B**), and > 83% of cells are insulin-producing cells (Q1 and Q2, **Figure S5B**). SC-islets were fixed with 4% PFA at room temperature for 30 min and embedded for cryosection. Section slides immunofluorescence staining corresponded with flow cytometry results, showing that the majority of the cells are CHGA and human C-peptide-positive cells (**Figure S5C**).

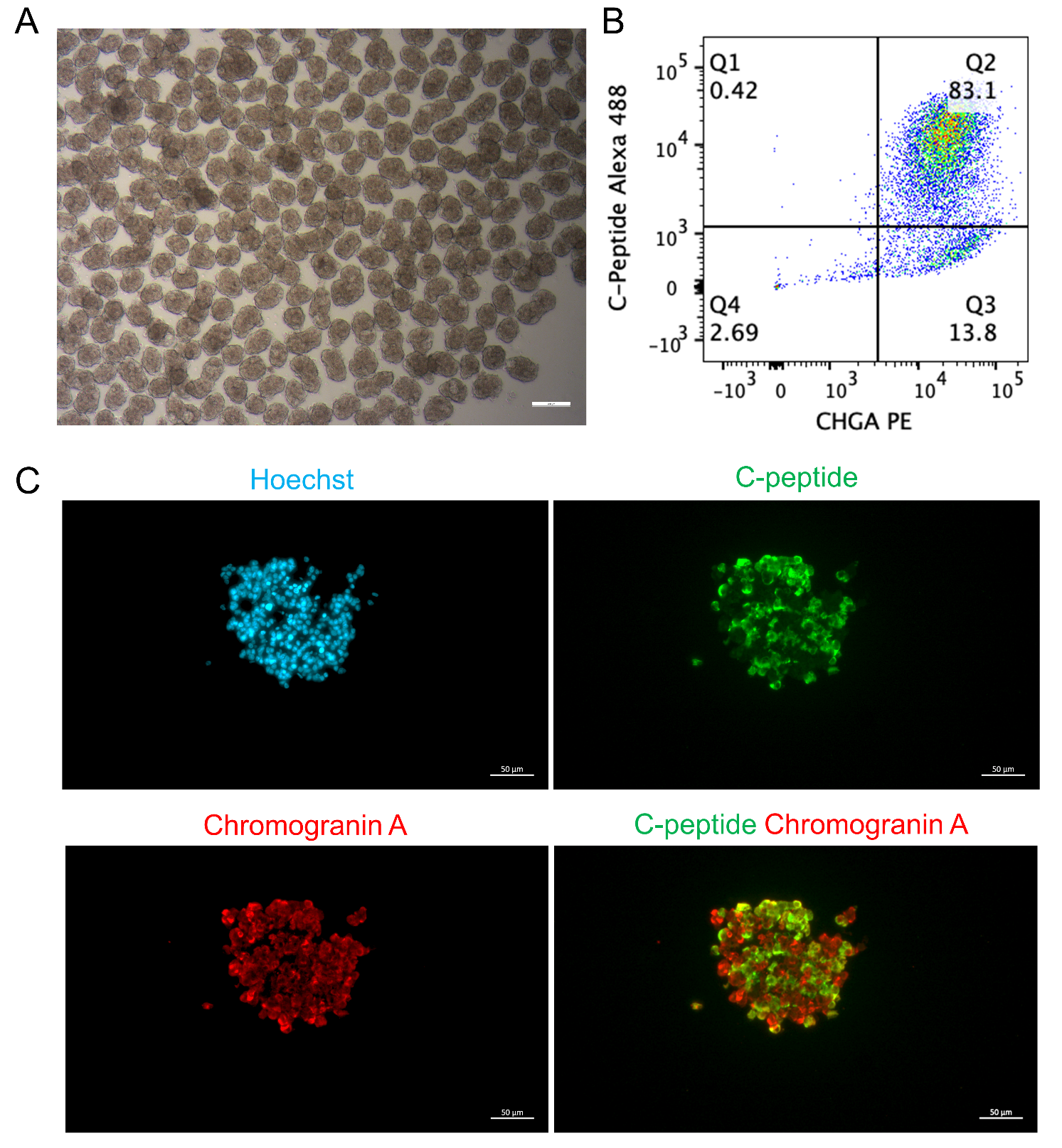

**Figure S5:** (**A**) Bright filed image of SC-islets. Scale bar = 200 µm. (**B**) Representative flow cytometry quantification (%) of dispersed SC-islets stained for C-peptide and Chromogranin A (CHGA). (**C**) Representative immunostaining images of SC-islets stained for Chromogranin A (red), C-peptide (green), and nuclei DNA with Hoechst (blue). Scale bar = 50 µm.
